# Mechanochemical Decoupling of ATP Hydrolysis and RNA Translocation in SARS-CoV-2 nsp13 by the L405D Mutation

**DOI:** 10.64898/2026.05.18.726016

**Authors:** Elham Fazelpour, Priti Roy, Kole J. Frederick, Martin McCullagh

## Abstract

SARS-CoV-2 nonstructural protein 13 (nsp13) is a highly conserved helicase that couples ATP hydrolysis to RNA translocation through long-range allosteric communication between its ATPase and RNA-binding domains. In prior work, we identified L405 as a key regulator of interdomain motions and proposed that the L405D mutation would disrupt this coupling by perturbing conformational translocations required for translocation [J. Phys. Chem. B 2024 v128 492–503]. Subsequent experiments confirmed that L405D attenuates helicase activity while largely preserving ATPase activity, implicating a breakdown in ATP-to-RNA coupling [J. Biol. Chem. 2026 v302 111198]. Here, we provide a data-driven explanation for this decoupling by combining Gaussian accelerated molecular dynamics (GaMD) simulations with Shape-GMM clustering and linear discriminant analysis. Whereas wild-type nsp13 exhibits both conformational selection and induction, L405D collapses the conformational landscape to operate predominantly through selection, eliminating ATP-induced structural transitions required for efficient catalytic cycling. This loss of induction traps the ATP-binding pocket in a mid-open conformation, impairing product release and reducing ATP turnover, while simultaneously disrupting coordinated motif–RNA interactions required for inchworm translocation. These findings establish that mutation-induced reshaping of conformational ensembles can modulate access to reaction-competent states, providing a general framework for understanding how targeted mutations disrupt catalytic function through allosteric ensemble remodeling in motor proteins.

## Introduction

The SARS-CoV-2 nonstructural protein 13 (nsp13) helicase is an essential viral motor whose function depends on coupling ATP hydrolysis to directional RNA translocation during genome replication. Like other superfamily 1 (SF1) helicases,^1,2^ nsp13 uses the free energy of nucleotide hydrolysis to unwind RNA duplexes and translocate along single-stranded RNA, enabling replication and transcription of the viral genome.^3–5^ Because disruption of this mechanochemical coupling can stall viral replication without necessarily preventing nucleotide binding or hydrolysis, nsp13 has emerged as an attractive target for antiviral drug development.^6,7^ Structural studies have shown that nsp13 contains two RecA-like ATPase domains (1A and 2A), an RNA-binding 1B domain, a stalk domain, and an N-terminal zinc-binding domain (ZBD), with long-range communication between the ATP-binding pocket and the RNA-binding cleft governing productive translocation (Figure 1).^8–12^ Understanding how this allosteric coupling is regulated is therefore central to explaining helicase function and dysfunction.

**Figure 1:**
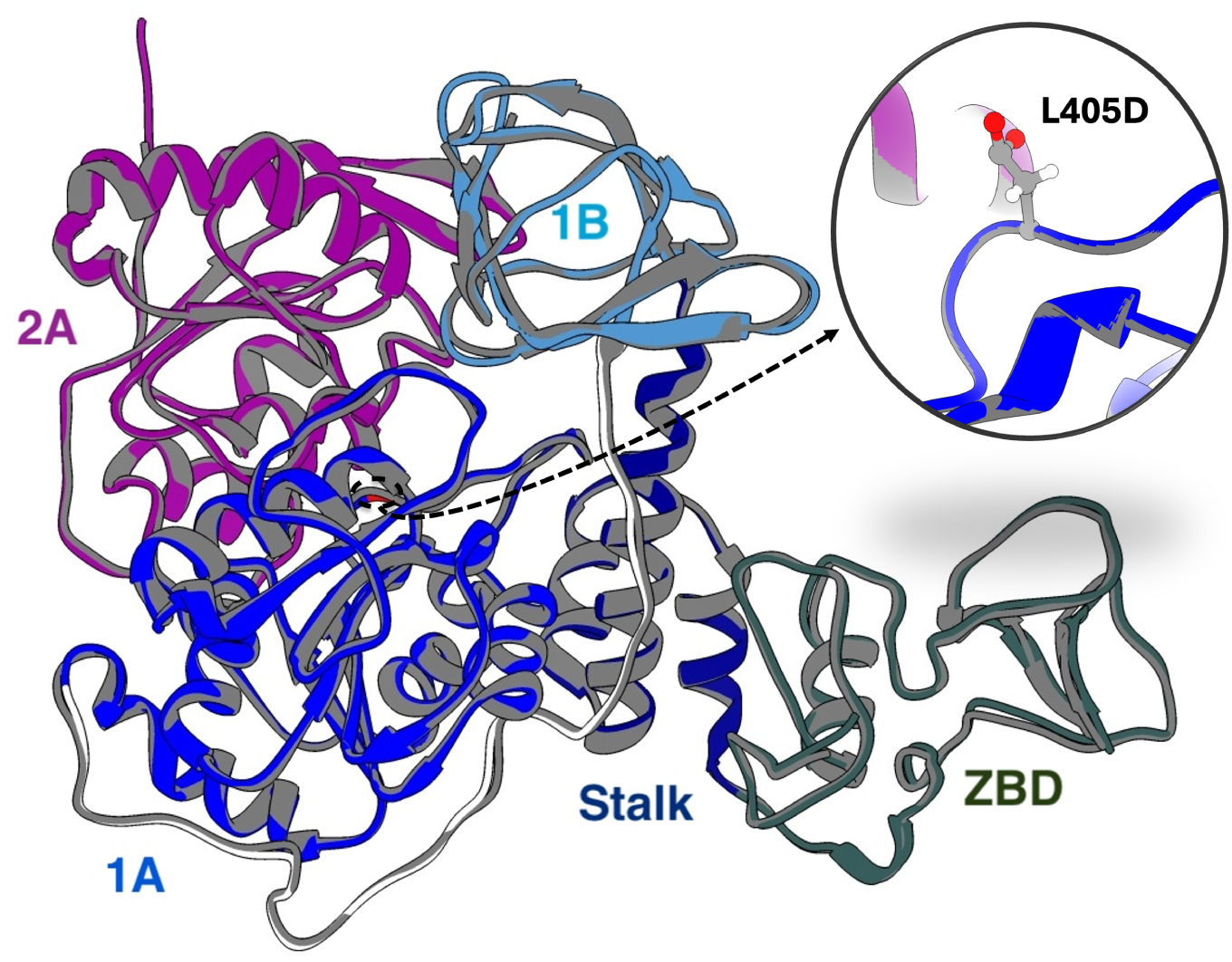
Depiction of SARS-CoV-2 nsp13 structure and substrate binding sites. Overlay of the L405D mutated nsp13 structure depicted with distinct subdomain colors and the SARS-CoV-2 structure (PDB: 7NN0^13^) in gray.

ATP-driven RNA translocation in helicases is governed by shifts in conformational ensembles, where ATP binding both selects pre-existing states and induces new reaction-competent conformations.^14–16^ In this framework, functional transitions arise through a balance between conformational selection and conformational induction that regulates interdomain cleft opening, motif–RNA contacts, product release, and directional stepping.^16–22^ Although nsp13 belongs to the SF1 helicase superfamily rather than SF2,^10^ it employs a related inchworm-like translocation mechanism in which coordinated binding and release of RNA by motifs Ia, IV, and V drive one-base movement along single-stranded RNA.^15,23–26^ Productive ATP hydrolysis therefore requires precise allosteric communication between the ATP-binding pocket formed by the 1A and 2A domains and the RNA-binding cleft formed by the 1A, 1B, and 2A domains. Mutations that perturb this communication can preserve ATP binding while disrupting mechanical translocation, leading to stalled or non-productive helicase cycles.

Previous computational and experimental studies identified L405 as a key allosteric regulator of ATP-dependent conformational transitions in nsp13.^16,27^ In our prior work, simulations along the ATP hydrolysis cycle showed that the presence of ATP does not substantially reorganize the ATP pocket itself, but instead shifts the relative positioning of the 1A, 1B, and 2A domains and alters the probability of RNA-cleft states associated with inchworm translocation.^14,16^ Linear discriminant analysis of these conformational changes identified residue L405, highlighted in the inset of Figure 1, as the strongest discriminator of open and closed interdomain states, indicating that it plays a central role in organizing the conformational ensemble required for productive ATP-driven translocation. Cryptic pocket analysis using PocketMiner^28^ further revealed that L405 is embedded within a transient, state-dependent pocket that emerges during the catalytic cycle (Figure S1), suggesting a regulatory role in stabilizing ATP-dependent structural transitions.^16^ Guided by these observations, we proposed the L405D mutation as a means to disrupt ATP-to-RNA coupling. Subsequent experiments confirmed that L405D strongly attenuates helicase activity while largely preserving ATPase function, demonstrating a breakdown in mechanochemical coupling rather than a loss of catalytic competence.^27^

The central unresolved question is how the L405D mutation selectively preserves ATP hydrolysis while disrupting ATP-dependent RNA translocation. We hypothesize that this decoupling arises because L405D reshapes the conformational ensemble of nsp13 by eliminating ATP-induced conformational induction and forcing the enzyme to operate almost entirely through conformational selection. Specifically, we propose that the mutation locks the dominant protein conformation in a mid-open ATP-pocket, closed RNA-cleft state (^P^C1) that persists regardless of nucleotide state, replacing the directional four-state wild-type progression with a restricted three-state ensemble that fails to couple nucleotide turnover to productive domain rearrangement. At the molecular level, we further propose that this conformational reshaping originates from perturbation of the native D534–R560 electrostatic network within Motif V, which decouples the ATPase site from the RNA-binding cleft and abolishes the coordinated motif–RNA register shifts required for productive inchworm translocation. To test this hypothesis, we performed Gaussian accelerated molecular dynamics (GaMD) simulations across four hydrolysis states (ssRNA, ssRNA+ATP, ssRNA+ADP+P*_i_*, and ssRNA+ADP), followed by Shape-GMM clustering and linear discriminant analysis to characterize mutation-dependent changes in protein and RNA-cleft conformational ensembles. These results provide a molecular mechanism for how mutation-induced ensemble remodeling decouples chemical and mechanical function in a viral motor protein.

## Methods

### Starting structure

To generate the mutated SARS-CoV-2 nsp13 helicase structure from the wild-type, we used crystallographic data (PDB: 7NN0^29^), with a resolution of 3.04 Å. This structure was co-crystallized with several components, including the substrate ANP, magnesium as a cofactor, water molecules, and zinc ions within the ZBD. Additionally, ssRNA was incorporated from an aligned RNA-bound Upf1 helicase structure (PDB: 2XZL^30^) as described in our prior study.^14^ We introduced the mutation by substituting LEU405 with ASP405, and applied this modification consistently across the substrate-bound states: ssRNA, ssRNA+ATP, ssRNA+ADP+P*_i_*, and ssRNA+ADP, in line with the wild-type simulation protocol.^16^

### System Preparation

For nsp13 and ssRNA, the force field parameters used were ff14SB^31^ and ff99bsc0*χ*OL3,^30,32^ respectively. ATP and ADP parameters were sourced from the AMBER parameter database.^33^ The P*_i_* parameter was taken from previous work by our group,^34^ and the three zinc ions in the nsp13 ZBD were parametrized using the MCPB module in AMBER.^35^ These zinc ions bond with residues in the following arrangements: Cys-Cys-Cys-Cys, Cys-Cys-Cys-HID, and Cys-Cys-HID-HIE in their respective pockets. Each substrate state was solvated using TIP3P water model with at least a 12 Å buffer, resulting approximately 207 K atoms after adding Na^+^ and Cl*^−^* ions to neutralize the charge and mimic a 0.1 M salt concentration.

### Simulation Details

All systems underwent energy minimization with a gradual removal of solute position restraints across 10 stages, each involving 2000 steps of steepest descent minimization. In the initial minimization step, a restraint of 500 kcal · mol*^−^*^1^ · Å*^−^*^2^ was applied to all atoms except water (excluding crystal water) and salt. The restraint was then progressively reduced to 50.0, 10.0, 1.0, and 0.1 kcal · mol*^−^*^1^ · Å*^−^*^2^ on the protein side chains, RNA, substrate, and Mg^2+^ ion over four steps. Subsequently, restraints on the protein backbone were gradually lowered in four phases, using 50.0, 10.0, 1.0, and 0.1 kcal · mol*^−^*^1^ · Å*^−^*^2^. The final minimization run was conducted without any restraints. Following minimization, protein complexes were heated incrementally from 0 to 300 K over 1 ns with a harmonic restraint of 40 kcal · mol*^−^*^1^ · Å*^−^*^2^ on protein, RNA, and substrate atoms. Following heating, six stages of pressure equilibration to 1 atm were conducted with progressively reduced harmonic restraints of 40.0, 20.0, 10.0, 5.0, 1.0, and 0.1 kcal · mol*^−^*^1^ · Å*^−^*^2^ on all protein and ligand atoms. The first equilibration stage was run for 1 ns, while the subsequent stages each ran for 200 ps. A restraint-free production run was then performed at constant temperature and pressure (*NPT*) for 10 ns, repeated in triplicate. In the ssRNA + ADP + P*_i_* substrate state, this step was extended to 500 ns to ensure that the Mg^2+^ ion’s octahedral coordination was conserved, particularly with SER289.^36^ From this production phase of conventional molecular dynamics (cMD), we recorded the minimum, average, maximum, and standard deviation values of the system’s potential energy. After the initial cMD simulations, we utilized Gaussian accelerated molecular dynamics (GaMD) simulations to improve conformational sampling across all systems. This approach included applying a boost to the entire system and dihedral potentials (igamd = 3), while keeping the other GaMD parameters at their default settings in Amber 18^37^ (*σ*_0P_ = 6 kcal · mol*^−^*^1^, *σ*_0D_ = 6 kcal · mol*^−^*^1^, *iE* = 1 or *E* = *V_max_*, nteb = 20,000,000, and ntave = 1,000,000 steps). Following the protocol, GaMD simulations commenced from equilibrated states obtained through brief cMD runs. All simulations were conducted using the GPU-accelerated AMBER18 package^37^ with a 2 fs integration time step, while hydrogen atoms were constrained with the SHAKE algorithm.^38^ Long-range electrostatic interactions were calculated using Particle Mesh Ewald, ^39^ applying a nonbonded interaction cutoff of 12 Å. Pressure and temperature were controlled using a Monte Carlo barostat and a Langevin thermostat, respectively. We conducted at least three GaMD simulations for each system, resulting in around 9 *µ*s per substrate states. In total, the ensembles accumulated to approximately 36 *µ*s.

### Shape-GMM Clustering

To quantify mutation-dependent changes in the conformational ensemble of nsp13 across the ATP hydrolysis cycle, we employed the size-and-shape space Gaussian mixture model (Shape-GMM) clustering algorithm.^40–42^ Shape-GMM provides a rigorous structural clustering framework that operates directly in aligned Cartesian coordinate space, allowing conformational states to be identified from collective domain motions without requiring predefined reaction coordinates or low-dimensional projections. This is particularly important for ATP-driven molecular motors such as nsp13, where function emerges from coordinated interdomain rearrangements rather than single geometric observables.

The effect of sampling on cluster populations is estimated by fitting 30 shapeGMM objects using different randomly selected training sets. Each training set is has equal weight from each substrate state. Following clustering, the cluster probabilities are reweighted for each hydrolysis state by using a second-order cumulant expansion of the boost potential, an energy-based correction method that effectively reduces the energetic noise introduced by the GaMD approach.^43^

### Linear Discriminant Analysis

Linear discriminant analysis (LDA) was used to identify the structural coordinates that most strongly distinguish functionally relevant conformational states and to define low-dimensional reaction coordinates for ATP-dependent transitions in nsp13. Unlike unsupervised dimensionality reduction methods, LDA is a supervised approach that maximizes variance between predefined conformational clusters while minimizing variance within each cluster, making it particularly well suited for identifying the dominant collective motions associated with ATP hydrolysis, product release, and RNA translocation.

Using cluster assignments obtained from Shape-GMM, aligned atomic coordinates were projected onto C − 1 discriminant vectors (where C is the number of conformational clusters) that optimally separate the states.^44^ These discriminant vectors provide an interpretable approximation to the reaction coordinate by highlighting the domain motions and residue-level displacements most strongly associated with transitions between clusters. In particular, the magnitude of individual residue contributions along the discriminant vectors was used to identify residues that act as allosteric regulators of conformational change. In our previous work, this analysis identified L405 as the strongest contributor to ATP-dependent opening and closing motions between the 1A, 1B, and 2A domains, motivating its selection for mutational analysis in the present study.

LDA was performed using the `scikit-learn`^45^ Python library with the singular value decomposition solver to ensure numerical stability in the high-dimensional coordinate space. The resulting discriminant projections were used both to assess cluster separability and to establish the mechanistic relationship between local residue perturbations and global conformational transitions.

### Model Corroboration

#### Presence of SF1 Helicase-like Conserved Contacts in the ATP Pocket and the RNA Cleft

In our current models, the ssRNA is derived from the crystal structure of the Upf1 helicase, while all substrates are produced through slight modifications of ANP, which is co-crystallized with SARS-CoV-2 nsp13.^29^ Given these modifications, it is crucial to assess the stability of these insertions by comparing protein contacts with those observed in crystal structures of other SF1 helicases. The contact between the SARS-CoV-2 nsp13 protein and its bound ligands including ssRNA, ssRNA+ATP, ssRNA+ADP+P*_i_*, and ssRNA+ADP systems are compared with analogous contacts observed in other SF1 helicases, such as Upf1^30^ and IGHBMP,^46^ to demonstrate that the selected starting structures are appropriate and remain stable during the simulations. A contact is considered to occur when a protein residue lies within 5 Å of ATP, ADP + P*_i_*, or ADP. The identities of these residues, along with the likelihood of their interactions with the ligands, are shown in Table S1, focusing on residues from key functional motifs.

## Results and Discussion

We propose that the L405D mutation decouples ATP hydrolysis from productive RNA translocation by eliminating ATP-driven conformational induction and trapping nsp13 in non-productive conformational states. In wild-type helicase function, the inchworm stepping cycle couples ATP turnover to directional ssRNA movement through coordinated communication between the ATP-binding pocket and the RNA-binding cleft, such that ATP binding, hydrolysis, and product release drive synchronized motif–RNA interactions required for forward translocation.^27^ We hypothesize that replacing L405 with aspartate disrupts this mechanochemical coupling by reshaping the conformational ensemble of nsp13, restricting access to reaction-competent states across the hydrolysis cycle and impairing ATP product release. To test this model, we analyze four hydrolysis-state systems (ssRNA, ssRNA+ATP, ssRNA+ADP+P*_i_*, and ssRNA+ADP) using Gaussian accelerated molecular dynamics simulations followed by Shape-GMM clustering and linear discriminant analysis. By comparing mutant conformational clusters to the corresponding wild-type ensembles, we distinguish whether ATP-dependent transitions arise through conformational induction (newly accessed states) or conformational selection (redistribution among pre-existing states). We first examine how L405D disrupts the motif Ia/IV/V interactions required for inchworm RNA translocation, then show how the mutation eliminates ATP-driven conformational induction, traps the ATP-binding pocket in closed or mid-open conformations, and impairs product release. Together, these analyses provide a molecular mechanism for how a single allosteric mutation decouples ATPase chemistry from productive mechanical translocation.

### L405D mutation disrupts the inchworm mechanism of RNA translocation

L405D disrupts the coordinated motif–RNA interactions required for directional inchworm translocation in nsp13. In the wild-type helicase, motifs Ia, IV, and V within the RNA-binding cleft execute synchronized cycles of loose-to-tight RNA binding that maintain a characteristic two-phosphate separation between the leading contact (motif IV) and trailing contacts (motifs Ia and V), enabling one-base (5*^′^* → 3*^′^*) stepping per ATP hydrolysis cycle.^13,14,16^ This coordinated inchworm mechanism couples ATP turnover to productive RNA unwinding during viral replication.^47,48^ Because L405 is positioned adjacent to motif V at a critical allosteric hinge, the L405D mutation is expected to perturb both RNA-cleft geometry and communication between the ATP-binding pocket and the RNA-binding motifs. Rather than supporting a single ATP-synchronized translocation pathway, the mutant is hypothesized to populate persistent, non-productive RNA-binding states that abolish coordinated stepping.^27^

L405D expands the RNA-cleft ensemble into four distinct conformational states, including non-productive binding configurations absent in wild-type nsp13. To characterize these cleft rearrangements, clustering was performed using the C*_α_* positions of motifs Ia, IV, and V together with the phosphate atoms of the bound ssRNA (Table S2), with trajectories from all hydrolysis states combined prior to analysis. The optimal number of RNA-cleft conformational clusters (^R^C1–^R^C4) was determined to be four based on the plateau in log-likelihood improvement and the minimum in the second derivative of the training-set curve (Figure S2a). Linear discriminant analysis further showed clear separation among these clusters (Figure S2b): LD1 primarily distinguishes the apo-like ^R^C1 from the more advanced states ^R^C2 and ^R^C4, while LD2 and LD3 resolve the distinct positioning of ^R^C3 and ^R^C4. Compared to the three-state RNA-cleft ensemble observed in wild-type nsp13, the mutant therefore samples a broader and less coordinated conformational landscape, indicating that L405D introduces additional substates that compete with productive ATP-coupled translocation.

ATP turnover no longer drives a single productive RNA-cleft progression in the L405D mutant, but instead redistributes the enzyme among competing non-productive substates. In the ssRNA-bound state, the ensemble is dominated by the apo-like ^R^C1 cluster (0.8±0.3), with only minor populations of the remaining clusters, Figure 2a, establishing ^R^C1 as the principal nucleotide-free conformation. ATP binding redistributes the population across all four clusters and shifts the dominant population to ^R^C3 (0.5±0.2), with secondary occupancy of ^R^C4 (0.3±0.2) and ^R^C2 (0.2±0.2), indicating that nucleotide binding drives a partial register shift rather than stabilizing a unique translocation-ready state. In the ADP+P*_i_*state, the ensemble returns to ^R^C1 (0.6±0.2) as the dominant cluster, with partial occupancy of ^R^C3 (0.2±0.1) and ^R^C4 (0.1±0.2), indicating that hydrolysis reverses rather than advances the RNA-cleft progression. Following the phosphate release, the ADP-bound state shifts toward ^R^C2 (0.7±0.1) and ^R^C1 (0.2±0.1), rather than progressing to a distinct post-hydrolysis conformation poised for reset. Unlike wild-type nsp13, where ATP binding and hydrolysis drive a coordinated and directional progression of RNA-cleft states, the L405D mutant repeatedly revisits overlapping conformations without establishing a single forward-translocation pathway. This loss of ATP-synchronized cleft progression explains how nucleotide turnover becomes decoupled from productive RNA movement.

**Figure 2:**
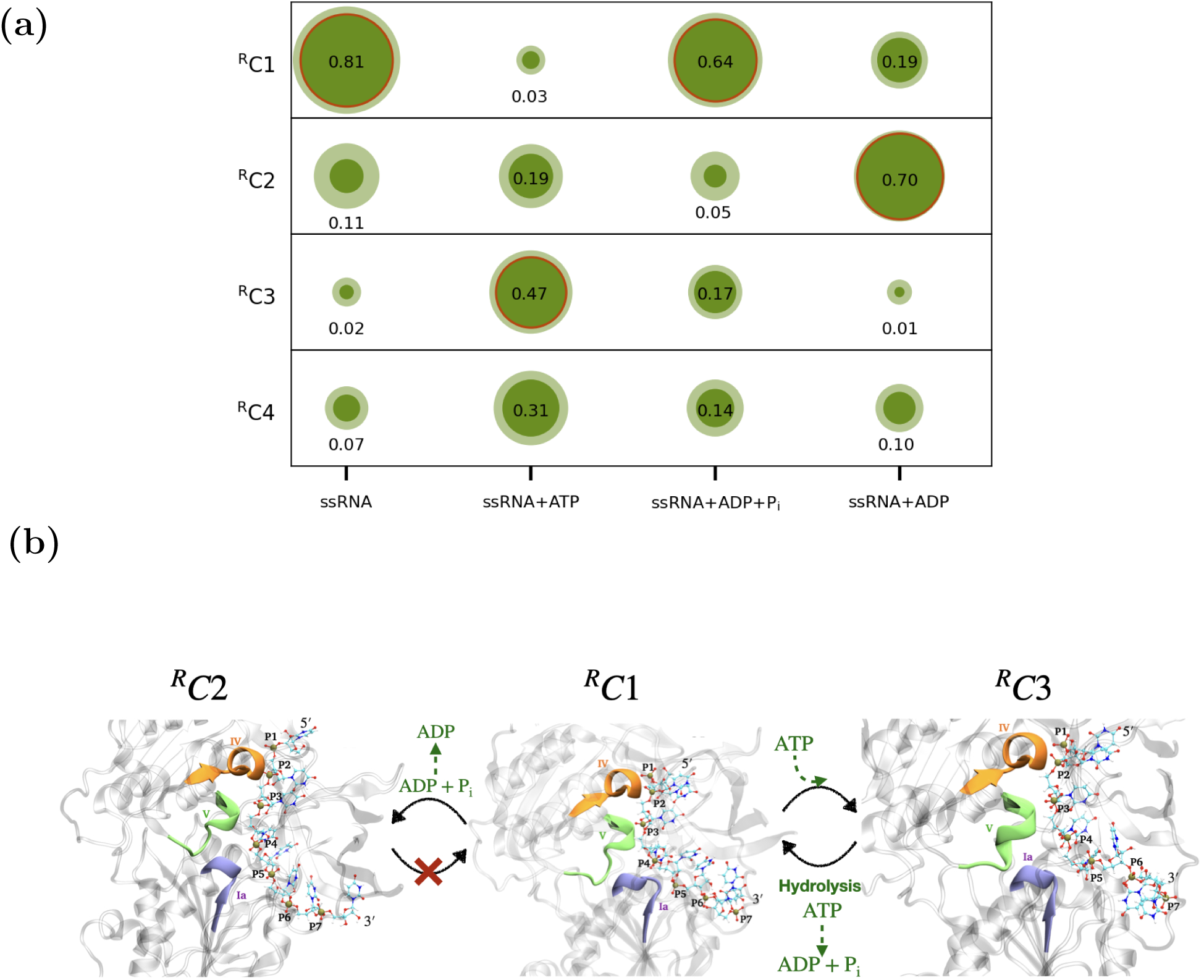
RNA cleft clustering and proposed disruption of directed translocation in the L405D mutant of nsp13. Clusters are identified by Shape-GMM clustering on the C*_α_* positions of residues in motifs Ia, IV, and V and ssRNA phosphates. (a) Reweighted probability of RNA cleft clusters explored by each system, presented in scatter form, where the size of each filled circle is proportional to the respective cluster’s probability and the surrounding halo represents the associated uncertainty estimated by the 2nd-order cumulant reweighting.(b) Representative structures of each RNA cleft cluster identified by the frame closest to the mean. Arrows indicate the futile ^R^C1 → ^R^C3 → ^R^C1 → ^R^C2 cycle imposed by L405D, in which ATP-driven leading-domain advancement (^R^C1 → ^R^C3) is reversed upon hydrolysis rather than consolidated, and ADP release is stalled by a tight-grip dead-end conformation (^R^C2), collectively uncoupling nucleotide turnover from productive inchworm translocation.

L405D disrupts the canonical two-phosphate separation required for directional inchworm stepping. In productive wild-type translocation, motif IV serves as the leading RNA contact while motifs Ia and V act as trailing contacts, maintaining a characteristic two-phosphate gap that enables one-base forward movement during each ATP hydrolysis cycle. ^16^ As shown in Tables 1 and 2, which contrast the motif–RNA contacts across all four conformational clusters of the L405D mutant, this coordination is systematically disrupted.

**Table 1:** Reweighted average separation distance (Standard Error) values between Motif IV, Ia, and V and the nearest ssRNA Phosphates (P) for clusters ^R^C1, ^R^C2, ^R^C3, and ^R^C4.

| residue | Average distance |  |  |  |
| --- | --- | --- | --- | --- |
|  | <sup>R</sup>C1(%) | <sup>R</sup>C2(%) | <sup>R</sup>C3(%) | <sup>R</sup>C4(%) |
| IV-P | 5.0(5) | 4.6(1) | 7.1(5) | 4.8(6) |
| V-P | 7.6(2) | 5.3(3) | 5.7(3) | 7.7(5) |
| Ia-P | 8.5(3) | 5.2(4) | 5.4(3) | 7.9(9) |

**Table 2:** Motif-Specific Closest Phosphates Reweighted Probability in Each RNA Cleft Clusters. The normalization was performed for each motif in each cluster.

| residue | Probability |  |  |  |
| --- | --- | --- | --- | --- |
|  | <sup>R</sup>C1(%) | <sup>R</sup>C2(%) | <sup>R</sup>C3(%) | <sup>R</sup>C4(%) |
| IV-P2 | 97 | 100 | 100 | 99.9 |
| IV-P3 | 3 |  |  | 0.1 |
| V-P3 | 100 | 100 | 100 | 99.9 |
| V-P4 |  |  |  | 0.1 |
| Ia-P4 | 2.1 | 99.7 | 100 | 99.9 |
| Ia-P5 | 97.9 | 0.3 |  | 0.1 |

Motif IV remains tightly associated with phosphate P2 across nearly all clusters, but the positioning of motifs Ia and V becomes uncoupled from this register, the precise motif-to-phosphate alignment required for directional stepping. In ^R^C1, the dominant ssRNA and ADP+P*_i_* state, motif IV contacts P2 tightly (5.0(5) Å), while motifs V and Ia remain relatively distant (7.6(2) and 8.5(3) Å, respectively), with motif Ia predominantly contacting P5 (97.9%), consistent with a disengaged pre-translocation state. Critically, the return to ^R^C1 upon hydrolysis – rather than advancement to a new register – means that the energy of ATP cleavage is dissipated into RNA release rather than forward stepping. The transition to ^R^C3, the dominant ATP-bound state, tightens motifs V and Ia (5.7(3) and 5.4 (3) Å) while motif IV loosens (7.1(5) Å), with Ia shifting to contact P4 exclusively (100%). This partial register advance is consistent with the leading domain stepping forward upon ATP binding, as expected in the first half of an inchworm stroke, this forward movement is not consolidated into net translocation. In ^R^C2, the dominant ADP-bound state, all three motifs simultaneously adopt their tightest RNA contacts across the entire cycle (4.6(1)– 5.3(3) Å), with motif Ia re-engaging P4 (99.7%). This represents a kinetic dead end:the enzyme achieves maximal RNA grip while bound to a spent nucleotide, preventing both RNA release and ADP dissociation required for productive reset. In ^R^C4, motif IV retightens (4.8(6) Å) while motif V and Ia disengage (7.7(5) and 7.9(9) Å), recapitulating a register similar to ^R^C1 without completing a productive reset. Critically, the canonical two-phosphate gap between motif IV and Ia is not preserved, and no cluster supports the forward power-stroke register seen in wild-type. Collectively, these RNA-cleft states demonstrate that L405D introduces extraneous binding configurations that scramble the strict alteration of tight/loose contacts required for directional stepping. Instead of a single, ATP-synchronized translocation event per cycle, the mutant samples a futile ^R^C1 → ^R^C3 → ^R^C1 → ^R^C2 loop that dissipate mechanical energy without net RNA advance. This mechanistic disruption, in which the chemical energy of hydrolysis is consumed but not converted into directional motion, directly accounts for the experimentally observed near-complete loss of helicase (unwinding) actively while ATPase function remains intact,^27^ establishing the RNA cleft as the decisive allosteric hub that couples nucleotide chemistry to mechanical translocation. These findings suggest that stabilizing non-productive cleft conformations may represent a promising antiviral strategy.

### L405D reshapes the ATP-dependent conformational ensemble by eliminating conformational induction

Unlike wild-type nsp13, which relies on both conformational selection and ATP-induced conformational induction,^16,49,50^ the L405D mutant operates almost entirely through conformational selection. In the wild-type helicase, ATP binding and product release drive a directional progression through four dominant protein conformational clusters (^P^C1–^P^C4), where some states are shared across hydrolysis intermediates through conformational selection and others are newly accessed only upon ATP binding or product release through conformational induction.^16^ In particular, ATP binding induces sampling of the ^P^C3 cluster, which is absent in the nucleotide-free ensemble, while the ADP-bound state preferentially accesses a distinct ^P^C4 cluster that is not populated in the ATP-bound state. These induced conformations are essential for coupling nucleotide turnover to productive domain rearrangements required for ATP-pocket opening, product release, and RNA translocation. The behavior of the L405D mutant differs fundamentally from this mechanism, indicating that the mutation does not simply shift populations but instead alters the underlying mode of conformational regulation.

L405D reduces the conformational diversity accessible during the ATP hydrolysis cycle, collapsing the four-state wild-type landscape into three dominant states. To characterize mutation-dependent changes in the global conformational ensemble, clustering was performed using the C*_α_* positions of the 1A, 1B, and 2A domains across the combined ssRNA-, ATP-, ADP+P*_i_*-, and ADP-bound ensembles. The optimal number of mutant protein conformational clusters was determined to be three (^P^C1–^P^C3) based on the plateau in log-likelihood improvement and the minimum in the second derivative of the training-set curve (Figure S3), in contrast to the four-cluster landscape previously identified for wildtype nsp13. Linear discriminant analysis further shows that these mutant clusters are well separated but reorganized relative to wild type (Figure 3a): LD1 primarily separates ^P^C2 from the remaining ensemble, while LD2 distinguishes ^P^C1 from ^P^C3. This partitioning differs qualitatively from the wild-type system, where ATP-dependent sampling of four distinct clusters reflects a broader conformational repertoire. As shown in Figure 3c, the cluster representatives reveal that L405D samples a reduced conformational space, suggesting restricted access to states required for efficient progression through the ATP hydrolysis cycle.

**Figure 3:**
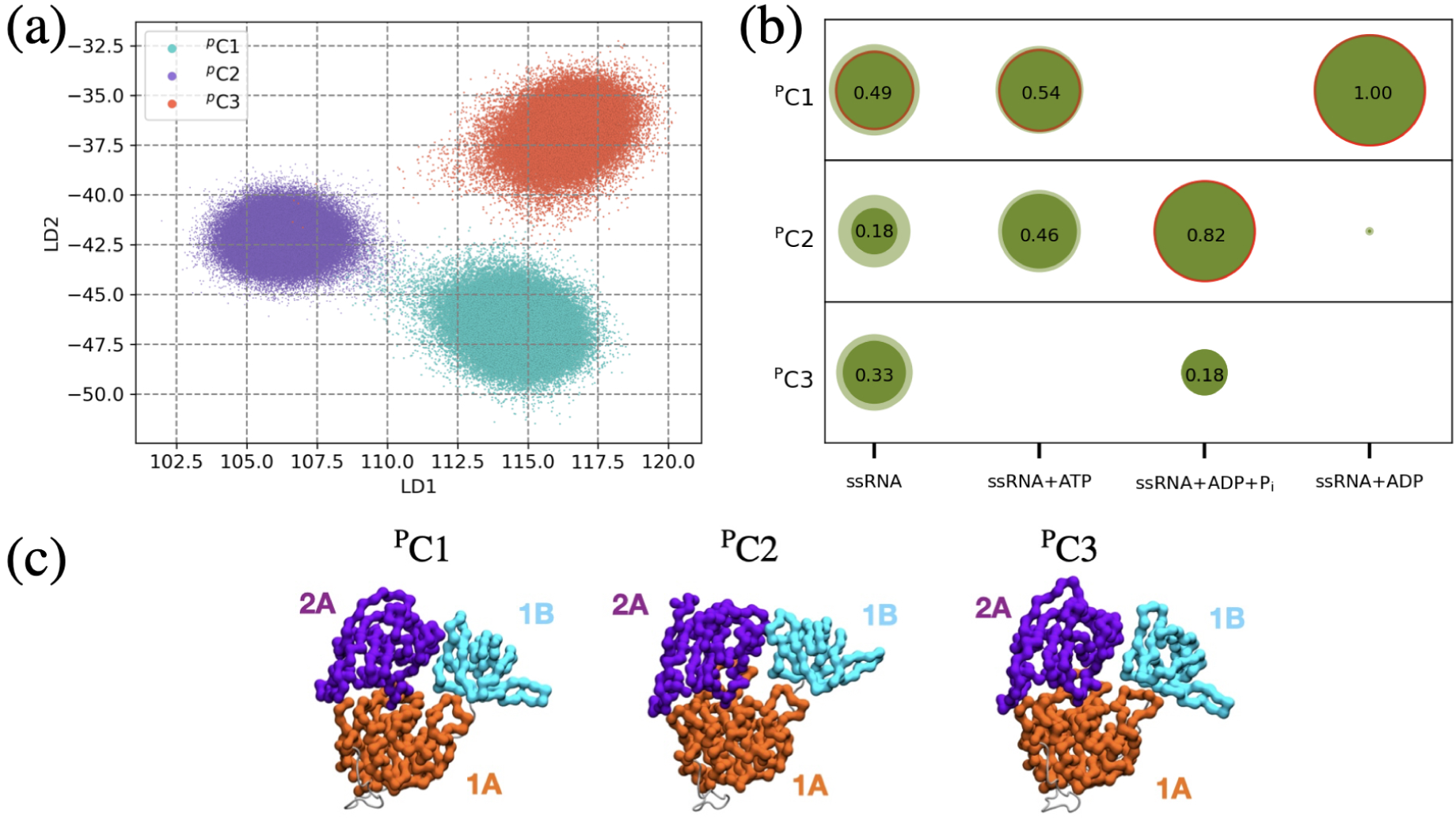
Full protein nsp13 conformational clusters across ligand-bound ensembles using Shape-GMM and LDA. (a) The 2D scatter plot denotes the projection of the amalgamated trajectory onto LD1 and LD2 eigenvectors determined using LDA on Shape-GMM clusters. (b) The reweighted probability of clusters explored by each system is plotted here and presented in scatter form. The size of each scatter is proportional to the respective cluster’s probability and the surrounding halo represents the associated uncertainty estimated by the 2nd-order cumulant reweighting. (c) Conformational clusters of nsp13 across ssRNA, ssRNA+ATP, ssRNA+ADP+P*_i_*, ssRNA+ADP ensembles, as determined by Shape-GMM clustering on the C*_α_* positions of domains 1B, 1A, and 2A.

No hydrolysis state in the L405D mutant accesses a uniquely induced conformation absent from the apo ensemble, indicating loss of ATP-driven conformational induction. In the ssRNA-bound state, the conformational ensemble is dominated by ^P^C1 (0.5±0.2) with substantial population of ^P^C3 (0.3±0.2), establishing these states as the principal nucleotide-free conformations (Figure 3b). ATP binding preserves ^P^C1 (0.5±0.1) as the dominant state while shifting secondary population toward ^P^C2, but critically does not introduce any cluster absent from apo ensemble, the hallmark signature of conformational induction is entirely absent. The ADP+P*_i_*–bound state produces the most pronounced redistribution, shifting the ensemble strongly toward ^P^C2 (0.8±0.0) with residual ^P^C3 (0.2±0.0), yet without accessing a new induced conformation. Following phosphate release, the ADP-bound state collapses entirely into ^P^C1 (1.0±0.0), returning to the same dominant nucleotide-free conformation rather than accessing a distinct post-hydrolysis state as seen in wild-type.^16^ The striking collapse of the ADP-bound state into ^P^C1, identical to the apo-dominant conformation, indicates that product release provides no conformational driving force for resetting the enzyme toward a new catalytic cycle. In contrast to wild-type nsp13, where ATP binding and product release induce sampling of conformations not present in the nucleotide-free state, every hydrolysis state of the L405D mutant remains confined to the same pre-existing set of clusters. These observations demonstrate that substrate binding drives only population redistribution among existing states, the hallmark of conformational selection, while the ATP-induced conformational induction required for productive catalytic cycling is lost.

The loss of conformational induction identifies L405 as a mechanistic pivot that couples ATP binding to productive domain rearrangement in nsp13. In the wild-type helicase, linear discriminant analysis of the protein conformational clusters showed that L405 contributes most strongly to the dominant ATP-dependent opening and closing motion between the 1A, 1B, and 2A domains, highlighting its central role in organizing the conformational response to substrate binding.^16^ Substitution of this residue with aspartate does not simply shift equilibrium populations within the existing ensemble; it eliminates the ability of ATP binding and product release to induce access to new reaction-competent conformations. The collapse of the ADP-bound state entirely into ^P^C1 is particularly telling: rather than the enzyme advancing to a new post-hydrolysis conformation that facilitates product release and reset, it reverts to the apo-like ground state with ADP still bound, consistent with impaired product release and futile cycling. As a result, substrate binding can no longer trigger the structural transitions required for ATP-pocket opening, coordinated RNA-cleft rearrangement, and efficient product release. The L405D mutation therefore converts a directional four-state catalytic cycle into a restricted three-state ensemble governed almost entirely by conformational selection.

The collapse of conformational induction predicts that ATP-pocket opening and product release should be kinetically impaired in the L405D mutant. In wild-type nsp13, ATP binding and hydrolysis drive access to induced conformations that facilitate coordinated opening and closing of the 1A–2A ATP-binding cleft, enabling efficient ATP binding, phosphate release, and eventual ADP dissociation. When this inductive component^21^ is lost, the enzyme becomes restricted to redistribution among pre-existing states, increasing the likelihood of trapping in closed or partially closed conformations that hinder product release. The complete collapse of the ADP-bound state into ^P^C1 further suggests that the conformational memory required to distinguish apo from product-bound states is erased, leaving the enzyme unable to couple ADP dissociation to the domain rearrangements needed to prime the next catalytic cycle. This provides a direct structural explanation for the experimentally observed attenuation of ATPase turnover^27^ despite preserved ATP binding and catalytic competence.^51^ We therefore next examine how mutation-dependent changes in interdomain cleft geometry alter accessibility of the ATP pocket and RNA-binding cleft across the hydrolysis cycle.

### Protein clustering reveals ATP-pocket closure and impaired product release

L405D traps the ATP-binding pocket in closed or mid-open conformations that impede product release and reduce catalytic turnover. Because ATP binding, phosphate release, and ADP dissociation are controlled by coordinated opening and closing of the interdomain clefts connecting the 1A, 1B, and 2A domains, changes in domain geometry provide a direct structural readout of catalytic accessibility. To quantify these motions, we analyzed the the reweighted probability distributions and mean values of the angles formed by vectors connecting the centers of mass of the three major domains (Figure 4a). The angle *θ*_1_ reports on the 1A–2A interdomain cleft that defines the ATP-binding pocket, while *θ*_2_ and *θ*_3_ describe the two RNA-cleft interfaces between 1A–1B and 2A–1B, respectively. Pairwise Welch’s *t*-tests confirmed that the three conformational clusters are statistically distinct across all three angle dimensions (*p <* 0.001 for most comparisons; see S2 Analyses and Figure S4 and S5 in the Supporting Information), supporting the conclusion that these clusters represent genuinely distinct conformational states rather than arbitrary partitions of a continuous distribution (Figure S4). Across the hydrolysis cycle, the dominant mutant clusters remain in ^P^C1 throughout the apo, ATP-bound, and ADP-bound states, with a transient excursion to ^P^C2 only in ADP+P*_i_*-bound state, indicating that the enzyme is persistently locked in an open ATP-pocket conformation that fails to couple nucleotide state to productive domain rearrangement (Figure 5).

**Figure 4:**
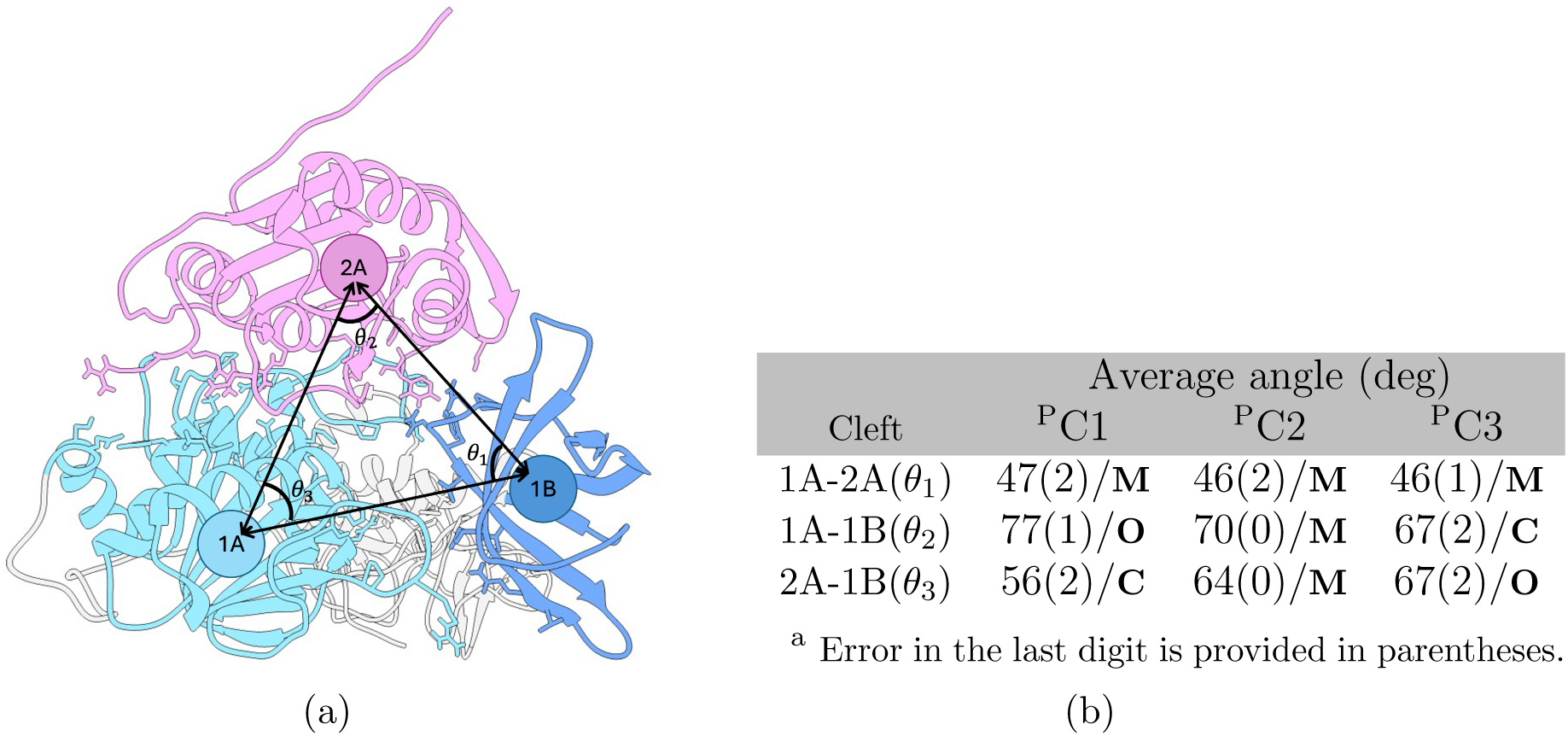
Interdomain angles within the nsp13 structure and the calculated reweighted average values in each protein cluster. (a) The spheres denote the centers of mass of the respective domains, and the black arrows represent the separation vectors between them. The arrow directions are defined relative to each specific angle calculation; for example, in computing *θ*_1_, the vectors are oriented from domain 1B toward domains 1A and 2A, respectively. (b) Reweighted average (standard error^a^) values of angles between subdomains for each protein cluster. **O** denotes ‘open‘, **M** denotes ‘mid-open‘, and **C** denotes ‘closed‘.

**Figure 5:**
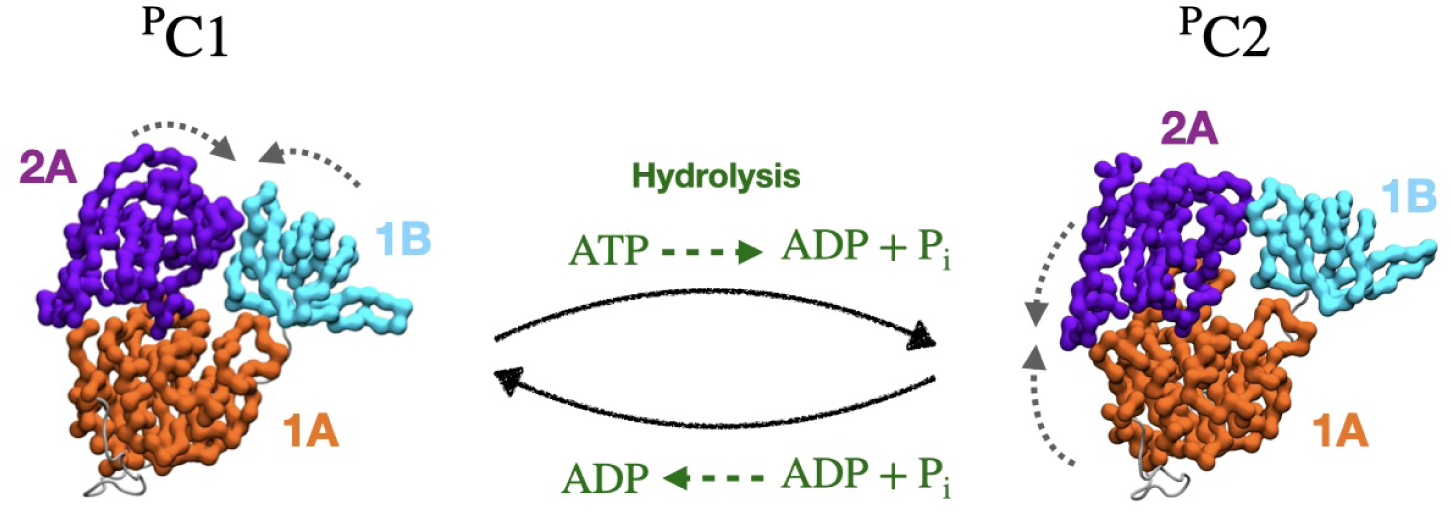
Proposed model of impaired translocation of L405D using representative structures for each protein cluster comprising the 1A (orange), 2A (purple), and 1B (cyan) domains of the L405D mutant. The protein clusters are organized in a cycle dominated by ^P^C1, with a transient excursion to ^P^C2 in the ADP+P*_i_*-bound state. The transitions between clusters, depicted by solid curved black arrows, are inferred from the sampled cluster populations during progression along the ATPase cycle: the ensemble remains ^P^C1 throughout the apo, ATP-bound, and ADP-bound states, shifting transiently to ^P^C2 only upon hydrolysis, before reverting to ^P^C1 upon ADP binding. The dotted gray curved arrows indicate intra-cluster domain motion guided by the interdomain cleft angle. To draw an analogy between this representation and the hydrolysis cycle, the entry of ATP and the release of ADP+P*_i_* and ADP are depicted in green.

L405D uncouples the ATP-binding cleft geometry from the nucleotide state, maintaining an open ATP pocket throughout most of the hydrolysis cycle and failing to drive the cleft closure required for catalytic progression. As shown by the hydrolysis-state cluster populations in Figure 3b, both the nucleotide-free ssRNA-bound ensemble and the ATP-bound ensemble are dominated by ^P^C1 (≈ 0.5). The corresponding interdomain geometries (Figure 4b) reveal the structural basis for this stalling. In the dominant ^P^C1 state, the the 1A–2A ATP-binding cleft is mid-open (*θ*_1_ = 47*^◦^*), the 1A–1B interface is open (*θ*_2_ = 77*^◦^*), and the 2A–1B RNA cleft is closed (*θ*_3_ = 56*^◦^*). The fact that ATP binding leaves the dominant cluster unchanged, with the ATP pocket remaining open, indicates that nucleotide binding fails to drive the cleft closure normally associated with catalytic engagement. The ADP+P*_i_*-bound state produces the only nucleotide-driven conformational transition in the cycle, shifting the ensemble strongly toward ^P^C2 (0.8±0.0), in which the ATP-binding cleft remains mid-open (*θ*_1_ = 46*^◦^*), the the 1A–1B interface adopts a mid-open conformation(*θ*_2_ = 70*^◦^*), and the 2A– 1B RNA cleft partially opens (*θ*_3_ = 64*^◦^*). However, this transition is reversed rather than consolidated upon phosphate release: the ADP-bound state collapses entirely back into ^P^C1 (1.0±0.0), the ATP-binding cleft remains mid-open (*θ*_1_ = 47*^◦^*) and re-closing the RNA-cleft (*θ*_3_ = 56*^◦^*) without advancing to a new post-hydrolysis register. This futile return to the apo- like ^P^C1 geometry with ADP still bound, an open ATP pocket incapable of driving product release, provides a structural explanation for impaired ADP dissociation and the experimentally observed reduction in ATP turnover.^27^ The 1A–2A cleft acts as the principal gate for ATP binding and product release, and L405D eliminates the nucleotide-dependent modulation of this gate. In the nucleotide-free state, the ATP-binding pocket is dominated by the open ^P^C1 (0.5±0.2) conformation with substantial population of the closed ^P^C3 (0.3±0.2), indicating that even in the absence of nucleotide the mutant enzyme samples both open and closed ATP-pocket geometries (Figure 4b). The narrow, well-defined probability distribution of *θ*_1_ for ^P^C1 (Figure S5, left panel) confirms that this cluster occupies a rigid conformational state with respect to the ATP-pocket geometry, in contrast to the broad, multimodal distributions observed for ^P^C3, which suggest a conformationally heterogeneous intermediate that interconverts between the more rigid ^P^C1 and ^P^C2 states. Upon ATP binding, the population remains anchored in ^P^C1 (0.5±0.1) while shifting secondary population toward ^P^C2, confirming that nucleotide binding fails to close the ATP pocket and does not stabilize a catalytically productive geometry. In the ADP+P*_i_*-bound state the ensemble shifts strongly to ^P^C2 (0.8±0.0) with residual ^P^C3 (0.2±0.0), representing the only stage of the cycle where cleft closure is observed, yet this occurs only after hydrolysis rather than before it, inverting the expected sequence of conformational events. After phosphate release, the ADP-bound state collapses entirely into ^P^C1 (1.0), reopening the ATP-binding cleft and returning to the same open-pocket geometry that dominates the apo ensemble. This inability to distinguish apo from ADP-bound conformations at the ATP pocket erases the conformational driving force required to couple ADP dissociation to resetting of the catalytic machinery, explaining how ATPase chemistry can remain accessible while overall ATP turnover is substantially reduced.^27^

While the ATP-binding pocket remains predominantly open and nucleotide-insensitive, the RNA clefts continue to sample open and closed geometries, but without the coordinated timing required for productive translocation. In the nucleotide-free and ATP-bound states, both dominated by ^P^C1, the the 2A–1B RNA cleft is closed (*θ*_3_ = 56*^◦^*) and the 1A–1B interface is open (*θ*_2_= 77*^◦^*), consistent with an RNA-engaged pre-translocation geometry. The absence of any ATP-driven conformational change means that the mechanical stroke normally associated with ATP binding is entirely absent. The ADP+P*_i_*-bound state shifts the ensemble to ^P^C2, where the 2A–1B cleft partially opens (*θ*_3_ = 64*^◦^*) and the 1A–1B interface moves toward mid-closure (*θ*_2_ = 70*^◦^*), suggesting that hydrolysis and phosphate retention partially reorganize the RNA-binding geometry. However, this rearrangement is not completed: the ADP-bound state immediately reverts to ^P^C1 with the 2A–1B cleft reclosed (*θ*_3_ = 56*^◦^*) and the 1A–1B interface reopened (*θ*_2_ = 77*^◦^*), without ever accessing the fully open RNA-cleft geometry (*θ*_3_ = 67*^◦^*) of ^P^C3. Because the only RNA-cleft rearrangement that occurs is confined to the ADP+P*_i_*-state and is reversed upon ADP binding rather than consolidated into a new translocation register, it cannot support the directional inchworm stepping required for productive RNA movement. The result is domain motion without efficient mechanical work.

To identify the specific residues driving these conformational differences, we decomposed the linear discriminant (LD) vectors from LDA into per-residue contributions and mapped their normalized magnitudes onto the protein structure (Figure S6). Along LD1, the highest-contributing residues are concentrated at the 1B–1A domain interface, with dominant peaks at residues 177–178 (domain 1B) and 308–309 (domain 1A), alongside a cluster of high-magnitude residues at 374–376 (domain 1A) and residue 514 (domain 2A), consistent with coordinated 1A–1B and 1A–2A cleft rearrangements identified above. Along LD2, discriminating residues localize predominantly to the 1A and 2A domains, with the highest contributions at residue 177 (domain 1B), residues 283 and 375 at the 1A–2A interface, residues 443–444 at the 1A–2A domain boundary, and residues 485, 515, 556–567 within domain 2A, reflecting the independent RNA-cleft rearrangements observed across the hydrolysis cycle. Notably, the elevated LD magnitudes observed near the 1A–2A domain boundary (residues 443–444) and within domain 2A (residues 514–515) along both discriminants suggest that the conformational discrimination between clusters originates from a localized structural perturbation in the vicinity of Motif V that propagates outward to the inter-domain interfaces. To determine whether L405D stabilizes genuinely novel conformations or merely redistributes population among wild-type-accessible states, we projected both trajectories onto the mutant- and wild-type-derived LD axes and compared their reweighted probability distributions (Figure S6). The Jensen–Shannon divergence between the two ensembles is large along both mutant-derived axes (*JS*_LD1_ = 0.69, *JS*_LD2_ = 0.69) and the wild-type-derived axis (*JS* = 0.62), with near-zero distributional overlap coefficients (*OC <* 0.03) across all projections. JS divergence values above 0.5 indicate ensembles that are more different than alike, and the consistency of this result across both coordinate systems rules out a coordinate artifact. These findings demonstrate that L405D does not redistribute population among pre-existing wild-type states but instead stabilizes a conformational landscape that is largely inaccessible to the wild-type enzyme, providing independent quantitative support for the mechanochemical decoupling described above.

These long-range conformational defects originate from electrostatic rewiring of the Motif V salt-bridge network introduced by the L405D substitution. In our previous study and recent experimental work on Motif V,^16,27^ L405 was proposed to function as an allosteric regulator by positioning the D534–R560 salt bridge that couples the ATPase site to the RNA-binding cleft. Replacing L405 with aspartate introduces a new negatively charged residue adjacent to D534, generating electrostatic competition for interaction with R560 and perturbing this communication pathway. Our distance analysis supports this model (Table 3). In both the ^P^C1 and ^P^C2 clusters, D534 maintains close contact with R560 (4.4–4.5 Å) while D405 remains more distant (6.5–6.8 Å), consistent with preferential stabilization of the native D534–R560 interaction throughout the accessible conformational ensemble. Importantly, neither cluster supports a configuration in which D405 competitively displaces D534 at R560; instead, the mutant residue remains peripherally positioned in both dominant states. This persistent native-like electrostatic arrangement, however, is insufficient to rescue productive conformational cycling, as the D405–R560 proximity is enough to perturb the precise interdomain communication required for nucleotide-dependent cleft modulation without fully redirecting the interaction network. By destabilizing a localized electrostatic hub, the L405D mutation propagates long-range effects on domain coordination, ATP-pocket accessibility, and RNA-cleft synchronization, providing the molecular origin of mechanochemical decoupling in nsp13.

**Table 3:**
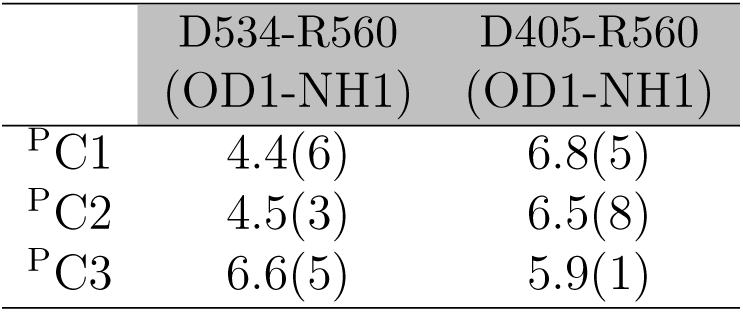
Average OD1–NH1 salt-bridge distances (Å) between D534–R560 and D405–R560 across predominant structural clusters (^P^C1 – ^P^C3). Values are reported as mean (standard deviation).

## Conclusions

This work provides a molecular mechanism for how the L405D mutation decouples ATP hydrolysis from productive RNA translocation in the SARS-CoV-2 nsp13 helicase. Although the mutant preserves ATP binding and catalytic competence, it abolishes helicase activity by disrupting the allosteric communication that links nucleotide turnover to directional inchworm stepping. At the protein-ensemble level, L405D eliminates ATP-induced conformational induction and collapses the four-state wild-type landscape into a restricted three-state ensemble. The dominant cluster ^P^C1, characterized by a mid-open ATP-binding pocket and closed RNA cleft, persists throughout the apo, ATP-bound, and ADP-bound states, with only a transient excursion to ^P^C2 during the ADP+P*_i_* state. This nucleotide-insensitive behavior means that ATP binding fails to drive catalytic engagement and ADP release fails to advance the enzyme into a new post-hydrolysis register.

At the RNA-cleft level, the mutation imposes a futile ^R^C1→^R^C3→^R^C1→^R^C2 cycle in which the forward register shift initiated upon ATP binding is reversed by hydrolysis rather than consolidated, and ADP binding locks the enzyme in a tight-grip dead-end conformation that prevents RNA release and productive reset. The canonical motif IV–Ia two-phosphate gap required for directional stepping is never preserved across the cycle. At the molecular level, these effects originate from perturbation of the native D534–R560 electrostatic network within Motif V, which propagates long-range disruption of ATP-pocket accessibility and RNA-cleft synchronization without abolishing catalytic activity.

Together, these findings establish L405 as a mechanistic pivot coupling ATP-driven conformational induction to directional translocation in nsp13, and identify the RNA-binding cleft as the decisive allosteric hub where chemical and mechanical cycles are coordinated. Stabilizing similarly non-productive cleft conformations may represent an effective antiviral strategy against SARS-CoV-2.

## Acknowledgments

The authors acknowledge the High Performance Computing Center at Oklahoma State University for providing computational resources supported in part through the National Science Foundation Grant OAC-1531128.

## Data Availability

The data that support the findings of this study, including simulation input files, analysis scripts, and representative trajectory data, are available from the corresponding author upon reasonable request.

