## Supporting Information for "Mechanochemical Decoupling of ATP Hydrolysis and RNA Translocation in SARS-CoV-2 nsp13 by the L405D Mutation"

#### S1 Supporting Table and Figures

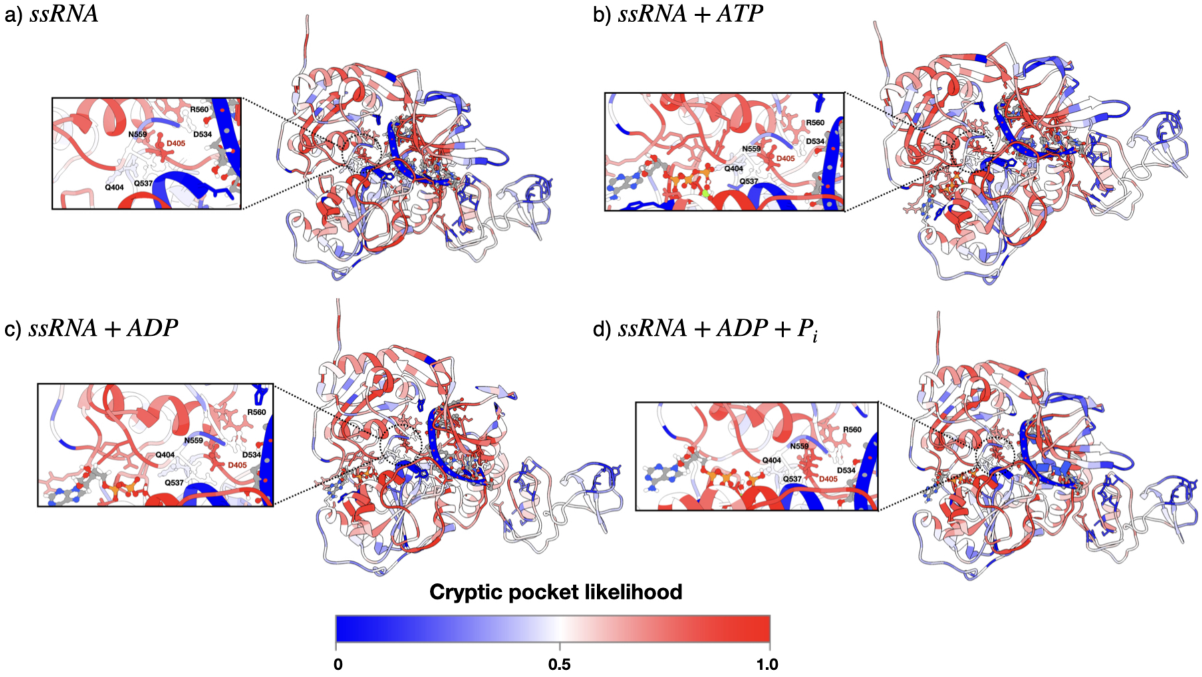


Figure S1: PocketMiner predictions on each substrate state. Proteins are colored from blue to red as predictions range from negative

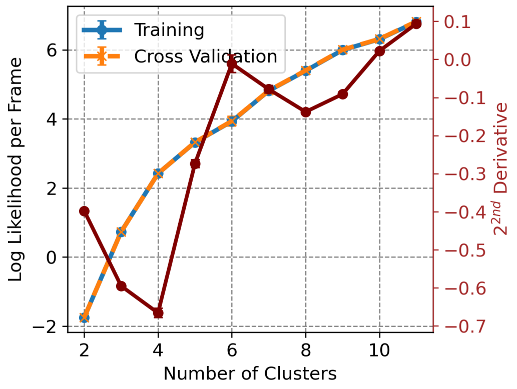


(a)

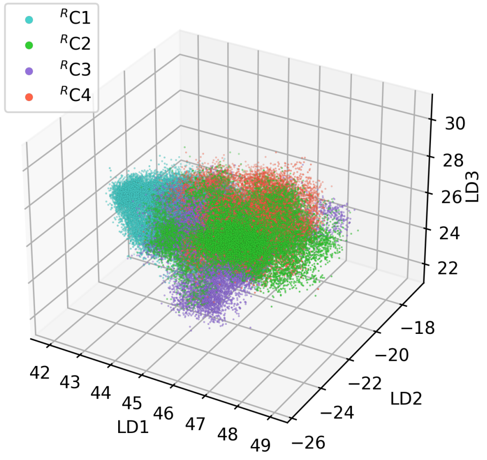


(b)

Figure S2: RNA cleft conformational clusters were determined by shape-GMM clustering. (a) Log likelihood per frame as a function of the number of clusters is plotted for the training (blue) and CV (orange) sets using the weighted Shape-GMM method. The change on slope of training set computed by the second derivative is plotted in red with values given on the right-hand $y$-axis. We used 80% of 3.6 million frames for training. (b) Projection of the amalgamated trajectory onto the three linear discriminants (LD1, LD2, and LD3) determined using LDA on Shape-GMM clusters.

### S2 Analyses

#### Inter-domain Cleft Angle Analysis

To characterize the conformational differences between the three protein clusters ($^P$C1, $^P$C2, $^P$C3) identified by shapeGMM, the inter-domain cleft angles $\theta_1$ (1A-2A), $\theta_2$ (1A-1B), and $\theta_3$ (2A-1B) were

Table S1: Motifs Ia, IV, and V residues in contacts with RNA "phosphates (P)" around 5.0 Å for SF1 RNA-bound helicases protein crystal structures and the percentage (%) of frames where the corresponding nsp13 residues are in contact with RNA(P) for each simulated substrate state.

| Motif | upf1 (2XZL) | IGHMBP2 (4B3G) | Nsp13 (7NN0) | ssRNA | ssRNA+ATP | ssRNA+ADP+$P_i$ | ssRNA+ADP |
|---|---|---|---|---|---|---|---|
| Ia | PRO460 | PRO243 | CYS309 | 2.5 | 55.8 | 36.4 | 17.4 |
| Ia | SER461 | SER244 | SER310 | 31.6 | 88.1 | 55.1 | 52.7 |
| Ia | ASN462 | ASN245 | HIS311 | 71.1 | 91.7 | 72.9 | 95.9 |
| Ia | VAL463 | ILE246 | ALA312 | 14.9 | 9.7 | 6.2 | 12.0 |
| IV | PRO731 | PRO540 | PRO514 | 41.8 | 24.5 | 19.4 | 16.5 |
| IV | TYR732 | TYR541 | TYR515 | 100 | 69.0 | 70.7 | 100 |
| IV | GLU733 | ASN542 | ASN516 | 100 | 47.8 | 63.6 | 100 |
| IV | GLY734 | LEU543 | SER517 | 68.2 | 30.3 | 35.6 | 86.1 |
| V | ALA764 | ASP565 | ASP534 | 59.0 | 96.4 | 97.9 | 73.5 |
| V | SER761 | SER563 | SER535 | 65.3 | 33.3 | 61.3 | 93.1 |

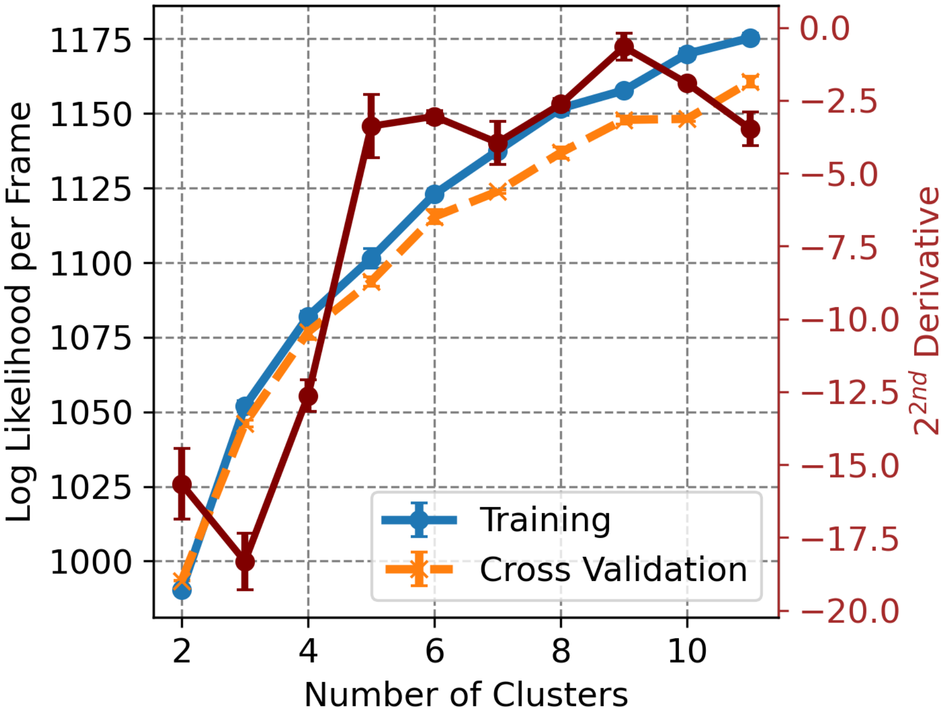


Figure S3: Identification of unique protein conformational clusters within the ensemble of nsp13-bound ssRNA, ATP+ATP, ssRNA+ADP+$P_i$, ssRNA+ADP, determined by Shape-GMM at the $C_\alpha$ resolution of domains 1B, 1A, and 2A. Log likelihood per frame as a function of the number of clusters is plotted for the training (blue) and CV (orange) sets using the weighted Shape-GMM method. The change on slope of training set computed by the second derivative is plotted in red with values given on the right-hand $y$-axis. We used 80% of ~300k frames for training.

Table S2: Residues of motifs in which $C_\alpha$ positions are used for distance calculations.

| Motif | Residue |
|---|---|
| Ia | HIS311 |
| IV | ASN516 |
| V | ASP534 |

computed for each frame of the trajectory subsampled every 10 frames. Each angle was defined as the angle subtended at one domain's center of mass by vectors pointing toward the centers of mass of the other two domains, computed using the dot product of the normalized inter-domain vectors:

$$\theta = \arccos\left(\frac{\mathbf{v}_1 \cdot \mathbf{v}_2}{|\mathbf{v}_1||\mathbf{v}_2|}\right) \tag{1}$$

The centers of mass were calculated for three protein domains: domain 1B (residues 151–226), domain 1A (residues 260–441), and domain 2A (residues 444–601), using only $C_\alpha$ atoms.

To account for the enhanced sampling bias introduced during the simulation, reweighted angle distributions were computed for each cluster independently. Frame weights were derived from the bias potential using the exponential reweighting scheme, where the statistical weight of frame $i$ is given by:

$$w_i = \frac{\exp(\beta V_{\text{bias},i})}{\sum_j \exp(\beta V_{\text{bias},j})} \tag{2}$$

where $\beta = (k_\text{B}T)^{-1}$, $T = 300$ K, and $k_\text{B} = 0.001987$ kcal mol$^{-1}$ K$^{-1}$. The reweighted mean angle $\langle\theta\rangle$ and standard error for each cluster were computed as:

$$\langle\theta\rangle = \sum_i w_i \theta_i \tag{3}$$

$$\sigma_{\langle\theta\rangle} = \sqrt{\frac{\sum_i w_i(\theta_i - \langle\theta\rangle)^2}{N}} \tag{4}$$

where $N$ is the number of frames in the cluster.

To assess whether the mean angles were statistically distinct between clusters, Welch's $t$-test[1] was applied to all pairwise cluster combinations ($^\text{P}$C1 vs $^\text{P}$C2, $^\text{P}$C1 vs $^\text{P}$C3, $^\text{P}$C2 vs $^\text{P}$C3) for each of the three inter-domain angles. Welch's $t$-test was chosen over Student's $t$-test because it does not assume equal variances between clusters, which is appropriate given the different population sizes and spread observed across the three conformational states. The test statistic is given by:

$$t = \frac{\bar{x}_1 - \bar{x}_2}{\sqrt{\frac{s_1^2}{n_1} + \frac{s_2^2}{n_2}}} \tag{5}$$

where $\bar{x}_1$ and $\bar{x}_2$ are the sample means, $s_1^2$ and $s_2^2$ are the sample variances, and $n_1$ and $n_2$ are the number of frames in each cluster. Statistical significance was defined at three levels: $p < 0.05$ (*), $p < 0.01$ (**), and $p < 0.001$ (***).

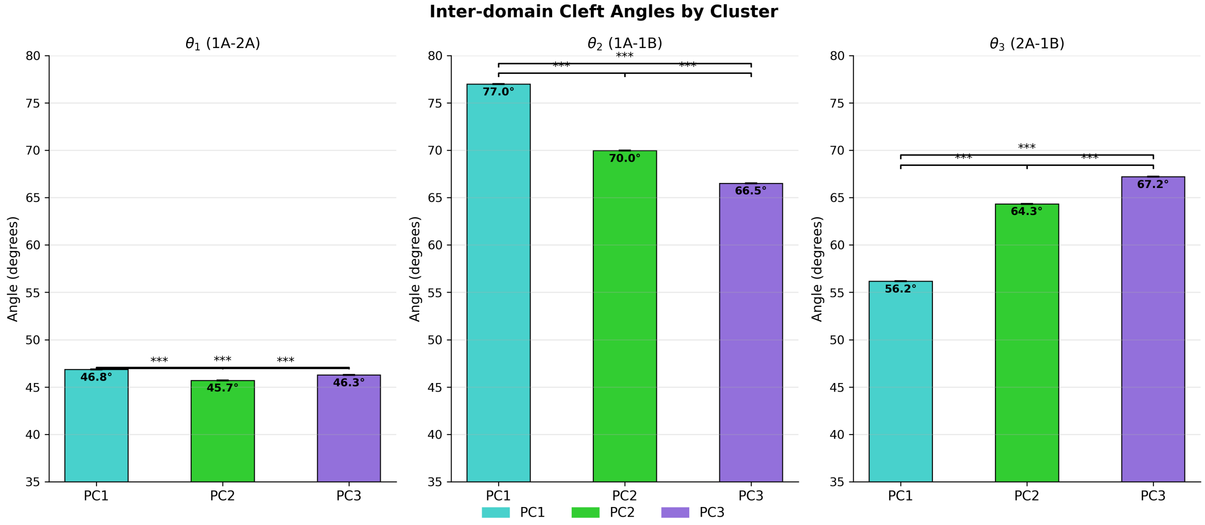


Figure S4: Reweighted mean inter-domain cleft angles for each conformational cluster with pairwise statistical significance. Bar plots show the reweighted mean values of (**Left**) $\theta_1$ (1A-2A cleft angle), (**Center**) $\theta_2$ (1A-1B cleft angle), and (**Right**) $\theta_3$ (2A-1B cleft angle) for clusters $^{\mathrm{P}}$C1 (medium turquoise), $^{\mathrm{P}}$C2 (lime green), and $^{\mathrm{P}}$C3 (medium purple). Error bars represent the reweighted standard error of the mean. Pairwise statistical significance between clusters was assessed using Welch's $t$-test, which does not assume equal variances between groups. Significance brackets indicate: ***$p < 0.001$, **$p < 0.01$, *$p < 0.05$, and ns (not significant) $p \geq 0.05$.

#### Reweighted Angle Probability Distributions

To visualize the conformational heterogeneity of each cluster with respect to the inter-domain cleft angles, reweighted probability distributions of $\theta_1$, $\theta_2$, and $\theta_3$ were computed independently for each conformational cluster ($^{\mathrm{P}}$C1, $^{\mathrm{P}}$C2, $^{\mathrm{P}}$C3). For each cluster, the angle values were histogrammed into 50 equally spaced bins over the observed range, with each frame contributing its statistical weight $w_i$ to the corresponding bin:

$$P(\theta_k) = \sum_{i \in \text{bin}_k} w_i \tag{6}$$

where the resulting distribution was normalized such that $\sum_k P(\theta_k) = 1$, yielding a reweighted probability distribution for each angle within each cluster.

The reweighted mean angle $\langle\theta\rangle$ for each cluster was computed as the weighted average:

$$\langle\theta\rangle = \sum_i w_i \theta_i \tag{7}$$

and is indicated in the distribution plots by a vertical dashed line. The width of each distribution reflects the conformational flexibility of the corresponding cluster along that inter-domain degree of freedom. Narrow distributions indicate conformationally rigid states, while broad or multimodal distributions indicate greater flexibility or the presence of multiple sub-states within a cluster.

The probability distributions for all three clusters were overlaid on the same panel for each angle to facilitate direct comparison of their positions and widths. Clusters $^{\mathrm{P}}$C1, $^{\mathrm{P}}$C2, and $^{\mathrm{P}}$C3 are represented in medium turquoise, lime green, and medium purple, respectively.

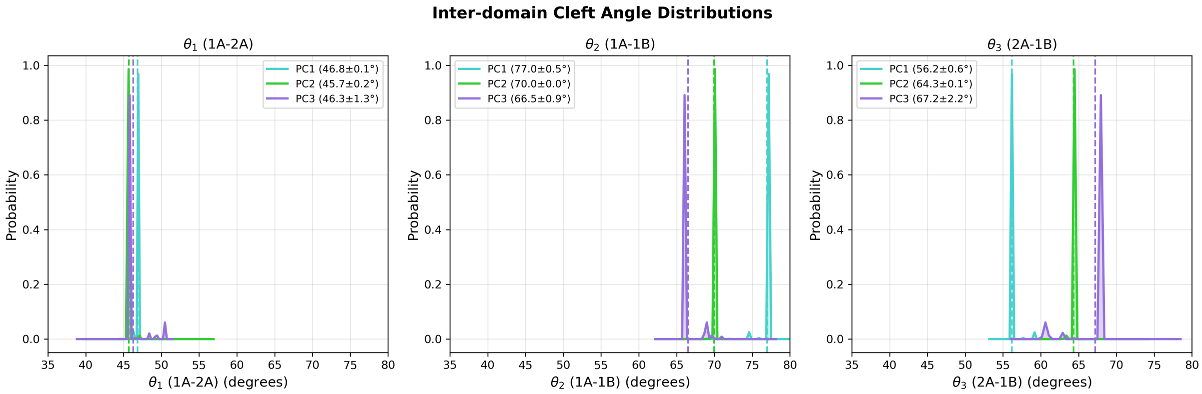


Figure S5: Reweighted probability distributions of inter-domain cleft angles for each conformational cluster. (**Left**) Distribution of $\theta_1$ (1A-2A cleft angle), (**Center**) distribution of $\theta_2$ (1A-1B cleft angle), and (**Right**) distribution of $\theta_3$ (2A-1B cleft angle) for clusters $^{\mathrm{P}}$C1 (medium turquoise), $^{\mathrm{P}}$C2 (lime green), and $^{\mathrm{P}}$C3 (medium purple). Distributions were computed by histogramming reweighted frame contributions into 50 equally spaced bins and normalizing to unit probability. Vertical dashed lines indicate the reweighted mean angle for each cluster. Narrow distributions reflect conformationally rigid states, while broad or multimodal distributions indicate greater flexibility or the presence of multiple sub-states within a cluster. Frame weights were derived from the bias potential using exponential reweighting at $T = 300$ K (see Methods).

#### Per-residue Linear Discriminant Contribution Analysis

To identify the specific residues driving the conformational differences between the three protein clusters ($^{P}$C1, $^{P}$C2, $^{P}$C3), the linear discriminant (LD) vectors obtained from LDA were decomposed into per-residue contributions. The LD vectors, of dimension $N_{\text{atoms}} \times 3$, where $N_{\text{atoms}} = 446$ C$_\alpha$ atoms (residues 151–596), were reshaped into a three-dimensional array of shape $(N_{\text{atoms}}, 3, N_{\text{LDs}})$, where the second dimension corresponds to the Cartesian components $(x, y, z)$ of each residue's contribution to each LD vector. The per-residue magnitude of contribution to each LD vector was then computed as the Euclidean norm of the Cartesian components:

$$m_{i,k} = \sqrt{x_{i,k}^2 + y_{i,k}^2 + z_{i,k}^2} \tag{8}$$

where $m_{i,k}$ is the magnitude of residue $i$'s contribution to the $k$-th linear discriminant, and $x_{i,k}$, $y_{i,k}$, $z_{i,k}$ are the corresponding Cartesian components of the LD vector for residue $i$.

To enable direct comparison of contributions across residues and between LD vectors, the per-residue magnitudes were normalized to the interval $[0, 1]$ using min-max normalization:

$$\tilde{m}_{i,k} = \frac{m_{i,k} - \min_j(m_{j,k})}{\max_j(m_{j,k}) - \min_j(m_{j,k})} \tag{9}$$

where $\tilde{m}_{i,k}$ is the normalized magnitude of residue $i$ along the $k$-th LD vector, and the minimum and maximum are taken over all $N_{\text{atoms}}$ residues. Residues with $\tilde{m}_{i,k}$ approaching unity are the primary structural determinants distinguishing the three conformational states along that LD axis, while residues with $\tilde{m}_{i,k}$ approaching zero remain relatively invariant across clusters.

The normalized per-residue LD magnitudes were visualized as bar plots as a function of residue index for LD1 and LD2 separately, with bars colored on a blue-to-red scale (coolwarm colormap) corresponding to low-to-high normalized magnitude. The top 10 residues with the highest normalized LD magnitude were identified and labeled for each LD vector to highlight the most discriminating regions of the protein. Additionally, the normalized magnitudes were mapped onto the representative structure of each conformational cluster as crystallographic B-factors to produce structure figures highlighting the spatial distribution of conformationally discriminating residues.

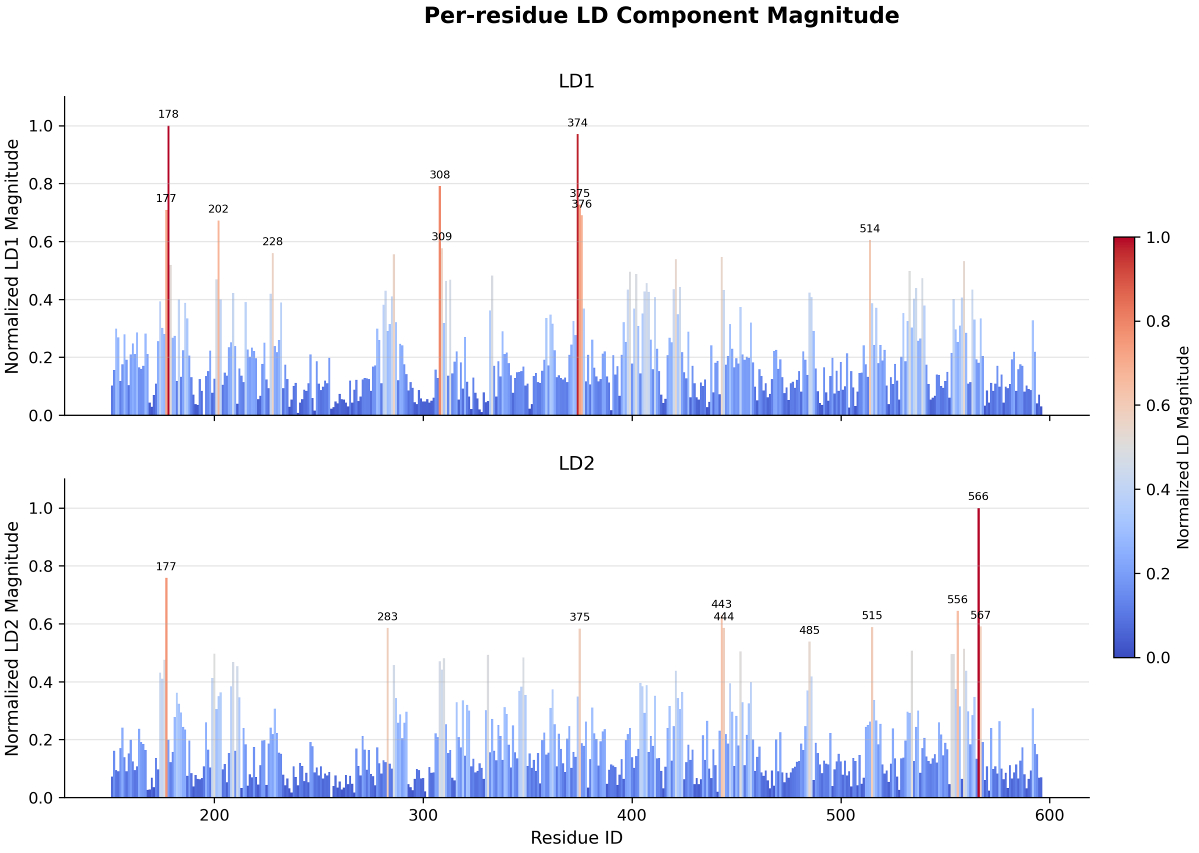


Figure S6: Per-residue contributions to the linear discriminant (LD) vectors separating the three conformational clusters. (**Top**) Normalized LD1 magnitude and (**Bottom**) normalized LD2 magnitude as a function of residue index (residues 151–596). The per-residue magnitude was computed as the Euclidean norm of the $(x, y, z)$ components of each residue's contribution to the LD vector, $\sqrt{x^2 + y^2 + z^2}$, and normalized to $[0, 1]$. Residues with high normalized magnitude (approaching 1, red) are the primary structural determinants distinguishing the three conformational states, while residues with low magnitude (approaching 0, blue) remain relatively invariant across clusters. The top 10 contributing residues are labeled for each LD vector.

#### Comparison of Mutant and Wild-type Conformational Ensembles

To assess whether the L405D mutant samples conformational states accessible to the wild-type enzyme, we compared the reweighted probability distributions of both trajectories projected onto the linear discriminant (LD) axes derived independently from each system. Mutant and wild-type $C_\alpha$ trajectories (residues 151–596) were flattened into feature vectors of dimension $N_{\text{atoms}} \times 3$ and projected onto the first two LD vectors derived from the mutant LDA model, as well as onto the first LD vector derived from the wild-type LDA model, yielding one-dimensional conformational projections for each system on each axis. Frame weights were computed independently for each trajectory using the exponential reweighting scheme described above.

The similarity between the mutant and wild-type reweighted distributions was quantified using two complementary metrics. First, the Jensen–Shannon (JS) divergence was computed as:

$$JS(P\|Q) = \frac{1}{2}D_{\text{KL}}(P\|M) + \frac{1}{2}D_{\text{KL}}(Q\|M) \tag{10}$$

where $P$ and $Q$ are the reweighted probability distributions of the mutant and wild-type projections, respectively, $M = \frac{1}{2}(P + Q)$ is their mixture distribution, and $D_{\text{KL}}$ denotes the Kullback–Leibler divergence. Both distributions were histogrammed into 50 equally spaced bins over the combined range of the two projections, normalized to unit probability, and regularized with a small epsilon ($\epsilon = 10^{-10}$) to avoid undefined logarithms. The JS divergence ranges from 0 (identical distributions) to 1 (completely non-overlapping distributions), with values above 0.5 indicating that the two ensembles are more different than alike.

Second, the overlap coefficient (OC) was computed as:

$$OC(P, Q) = \sum_k \min(P_k, Q_k) \tag{11}$$

where the sum runs over all histogram bins $k$. The overlap coefficient ranges from 0 (no shared conformational space) to 1 (identical distributions) and provides a direct measure of the fraction of conformational space sampled by both systems. Both metrics were computed on the mutant-derived LD1 and LD2 axes, as well as on the wild-type-derived LD1 axis, to ensure that the observed separation was not an artifact of the coordinate system used for projection.

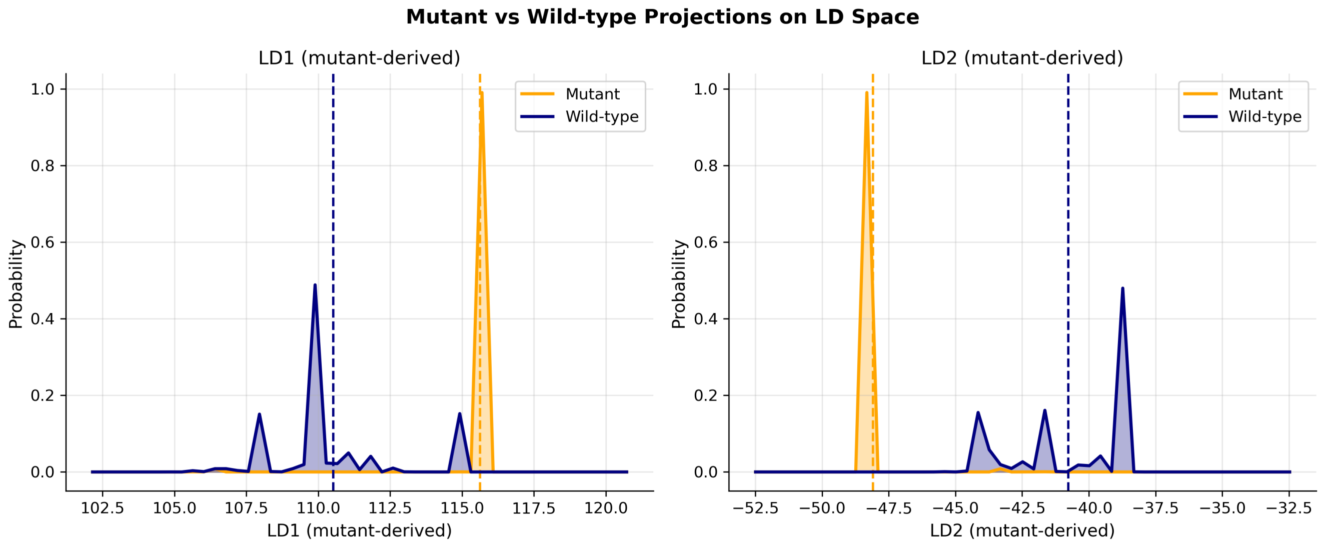


Figure S7: Comparison of mutant and wild-type conformational ensembles projected onto the mutant-derived linear discriminant axes. Reweighted probability distributions of the L405D mutant (orange) and wild-type (navy) trajectories projected onto (**Left**) LD1 and (**Right**) LD2, both derived from the mutant LDA model. Vertical dashed lines indicate the reweighted mean projection value for each system. The mutant distribution is narrow and highly localized, while the wild-type distribution is broader and multimodal, sampling multiple distinct regions along each axis. The near-complete absence of distributional overlap ($OC < 0.002$ for both axes) and large Jensen–Shannon divergence ($JS > 0.69$) confirm that the two systems occupy largely distinct regions of conformational space, indicating that L405D stabilizes conformations that are rarely, if at all, accessible to the wild-type enzyme.
